# Read correction for non-uniform coverages

**DOI:** 10.1101/673624

**Authors:** Camille Marchet, Yoann Dufresne, Antoine Limasset

## Abstract

Next generation sequencing produces large volumes of short sequences with broad applications. The noise due to sequencing errors led to the development of several correction methods. The main correction paradigm expects a high (from 30-40X) uniform coverage to correctly infer a reference set of subsequences from the reads, that are used for correction. In practice, most accurate methods use *k*-mer spectrum techniques to obtain a set of reference *k*-mers. However, when correcting NGS datasets that present an uneven coverage, such as RNA-seq data, this paradigm tends to mistake rare variants for errors. It may therefore discard or alter them using highly covered sequences, which leads to an information loss and may introduce bias. In this paper we present two new contributions in order to cope with this situation.

First, we show that starting from non-uniform sequencing coverages, a De Bruijn graph can be cleaned from most errors while preserving biological variability. Second, we demonstrate that reads can be efficiently corrected via local alignment on the cleaned De Bruijn graph paths. We implemented the described method in a tool dubbed BCT and evaluated its results on RNA-seq and metagenomic data. We show that the graph cleaning strategy combined with the mapping strategy leads to save more rare *k*-mers, resulting in a more conservative correction than previous methods. BCT is also capable to better take advantage of the signal of high depth datasets. We suggest that BCT, being scalable to large metagenomic datasets as well as correcting shallow single cell RNA-seq data, can be a general corrector for non-uniform data. Availability: BCT is open source and available at github.com/Malfoy/BCT under the Affero GPL License.

## Introduction

### Motivations

Fast and accurate pre-filters are key to many bioinformatics applications such as assembly (to remove spurious nucleotides and propose more contiguous assemblies), variant calling, genotyping or mapping (to help reduce the noise ratio and increase the power of these methods). Removing errors is also determinant to reduce the quantity of data to index, in particular for *k*-mer approaches. Thus, a good read correction should allow to index less, though more meaningful, data.

While numerous and very efficient methods exist for genomic correction, less methods were dedicated to short-reads datasets presenting uneven coverage^1^ such as transcriptomics or metagenomics. Short read sequencing (shotgun and single-cell) remain the widest way to access transcriptomics, metagenomics and metatranscriptomics data despite the growing applications of long reads (PacBio, Oxford Nanopore). Partly because of the relative lack of correction methods, most transcriptomic/metagenomic softwares include their own pre-filters, which is not in favor of homogeneous pipelines. This motivates efficient and reliable correction of non-uniform coverage short-read data.

## Background

### *k*-mer spectrum methods

These correction methods assume a uniform sequencing coverage. With the second assumption that sequencing errors are uniformly distributed along read sequences, then there must exist a coverage level for *k*-mers (the set of read substrings of length *k*) at which it becomes highly unlikely to observe errors. Thus, they compute distribution of the *k*-mers’ number of occurrences (spectrum) in the dataset, and rely on an abundance threshold that is chosen to distinguish “weak” (error-prone) *k*-mers from “solid” *k*-mers. “Weak” *k*-mers are then corrected by converting them to corresponding solid ones. Because they are conceptually simple and only rely on computationally efficient structures [15, 18, 28], these methods have been preferred to other approaches relying on suffix structures [25], [21], [13] and probabilistic models or multiple sequence alignment [23].

### Graph-based methods

Recently, methods based on De Bruijn graphs emerged for read correction. The rationale is that a set of *k*-mers can efficiently be represented in such a graph, and that a read can be corrected by its alignment on the graph that contain a low amount of errors. Following this idea, the two methods [22] [24] were specifically designed to correct long reads from PacBio or Oxford Nanopore technologies (therefore, they are out of the scope of short read correction, as shown in [17]). Lastly, Bcool [17] introduced a new approach dedicated to short reads. The approach first constructs a De Bruijn graph and applies heuristics to remove most of the erroneous *k*-mers from the graph. Thereafter, the reads are aligned to this cleaned graph which is used as a set of reference sequences to correct them. This allows to take into account more global information for the correction by relying on graph paths instead of *k*-mers.

### Adaptation to non-uniform coverage

In transcriptomics, data reflect the gene expression levels. Consequently, relative low-frequency *k*-mers are not always the mark of a sequencing error, as they may pertain to lowly expressed regions. Moreover, in a De Bruijn graph, alternative splicing or alternative transcription events produce several alternative correct *k*-mers in the same graph region. Those properties induce a harder correction process. In metagenomics, the mixture of unevenly represented genomes yields similar issues.

The first approach to face this issue was SEECER [14], combining a multiple sequence alignment to hidden Markov models, but was poorly scalable. BayesHammer and Rcorrector [27] both derive from the idea to compute groups of neighbors *k*-mers, from which “solid” *k*-mers are identified, and used to correct “weak” *k*-mers from the same group. BayesHammer (designed for single-cell datasets) proposes a clustering technique based on bayesian statistics derived from the k-means clustering technique to find groups of close *k*-mers according to the Hamming distance. Rcorrector is designed for transcriptomic datasets, and proposes to choose among sets of *k*-mers from possible alternative paths in the De Bruijn graph to correct an error. The path chosen must not contain *k*-mers counted less than a fixed threshold, and must not be incoherent with the read’s *k*-mer counts. Finally, to our knowledge, there exist no method specifically dedicated to the correction of shotgun metagenomics data.

### Contribution

In this paper, we show that important algorithmic novelties are required to propose an uneven coverage corrector based on De Bruijn graph cleaning and alignment. We demonstrate that:

- *k*-mer set cleaning steps, whether they are *k*-mer spectrum approaches, assembler pre-filters or Bcool, have room from improvement, and we present a new approach with better results,
- a graph approach can be combined with a likelihood model for scoring the unitigs according to their probability to be generated by errors. Such approach allows to save more “weak” although meaningful *k*-mers,
- local read alignment on the graph provides better results for read correction (more read corrected and less errors introduced in reads) than the current techniques.

## Methods

### Preliminaries

Let Σ be an alphabet of fixed size, in this work we assume Σ = {*A,C,G,T*}. A *k*-mer is a sequence *s*∈Σ_*k*_ (i.e of length |*s*| = *k*). We consider a read dataset as a multiset *R*⊂Σ* and an integer *k*, and define the De Bruijn graph as a directed graph *G*_*k*_(*R*)=(*N,E*) where *N* is the set of distinct *k*-mers that appear in *R*, and for *u,v*∈*N*,(*u,v*)∈*E* if and only if *u*[2,*k*]=*v*[1,*k*-1] (see (1) in Figure 1).

**Figure 1:**
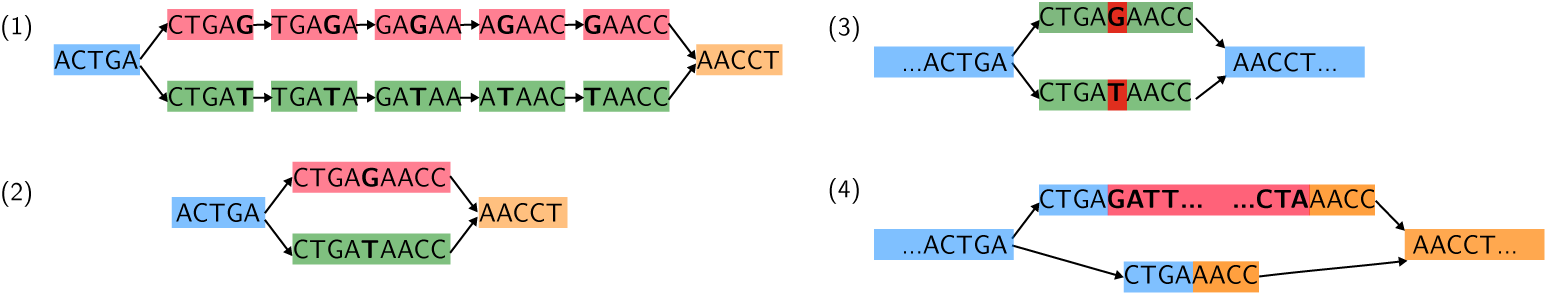
(1) De Bruijn graph with *k* = 5. Nodes are *k*-mers. (2) Same, compacted, De Bruijn graph. The *k*-mers constituting simple paths (red CTGAG,TGAGA,GAGAA,AGAAC,GAACC and green CTGAT,TGATA,GATAA,ATAAC,TAACC) have been merged by concatenating two consecutive *k*-mers while removing the overlapping part of the second one. Each node of this graph is called a unitig. (3) Effect of a single substitution in a compacted De Bruijn graph. The alternative versions of a nucleotide (red G/T) create two unitigs of the same lengths, that only differ by one position. Such an event can originate from a sequencing error (G is an error while T is genomic) or a SNV. (4) Expected topology in the case of an alternative splicing event in a transcriptomic De Bruijn graph (see [20] for a formalization). The red part is the sequence of an alternatively spliced exon or an insertion/deletion (indel). The inclusion/exclusion of exon-length or indel sequences creates patterns where branching nodes are connected to paths that may differ in length and sequence content.

The out-neighborhood (respectively in-neighborhood) of a node *u* is {*v*∈*N*|(*u,v*)∈*E*} = *N*^+^ (respectively {*v*∈*N*|(*v,u*)∈ *E}* = *N*^−^ and its in-degree *d*^+^(*u*) (respectively out-degree *d*^−^(*u*)) is |*N*^+^| (respectively |*N*^−^|). A node *u* is called a *branching node* if *d*^+^(*u*)>1 or *d*^−^(*u*)>1.

We call *simple path* in *G*_*k*_ a set of nodes *s* = *u*_0_,*…u*_*n*_ such that, for each 0≤*i*≤*n*, (*u*_*i*_,*ui*+1)∈*E, d*^+^(*u*_0_) = 1,*d*^−^(*u*_*n*_) = 1 and for each 1≤*i*≤*n*-1,*d*^+^(*u*_*i*_)=*d*^−^(*u*_*i*_)=1. A simple path *s* = *u*_0_,*…u*_*n*_ can be compacted by concatenating *u*_0_ to *u*_*i*_[*k*-1] for each 0*<i*≤*n* into a single node of length *k*+*n*-1, and by keeping only edges to in-neighbors of *u*_0_ and to out-neighbors of *u*_*n*_ (see (2) in Figure 1). We call a graph whose simple paths have been compacted a *compacted De Bruijn graph*. Resulting compacted nodes are called *unitigs*. In the following, we work with this node-centric, compacted version of a De Bruijn graph.

### Algorithm overview

We present an approach to correct reads by using a 2-steps procedure. First a De Bruijn graph is built fom the reads’ *k*-mers. This graph is iteratively cleaned (unitigs are removed, see section 2.3) using topological, textual and unitig coverage information (the unitig coverage being 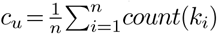 the *k*-mers of the unitig *u* of size *n*+*k-*1). Then, reads are mapped to the cleaned graph, using a procedure involving coverage estimation and local alignment, to decide which path is the most eligible to correct each read (see section 2.4).

### Graph cleaning

#### Graph construction and hard filters

To build a reference De Bruijn graph, we rely on Bcalm2 [5] to build a raw De Bruijn graph from all non-unique *k*-mers of the dataset. *k*-mers are extracted and counted, then Bcalm2 constructs the unitigs and computes their average coverage. Optionally, we can apply a filter on this coverage, removing low coverage unitigs (useful in case (5), Figure 2). While this kind of filtering has been shown to be efficient in genomics [17], such a threshold could suppress poorly represented but relevant *k*-mers. Unless said otherwise we did not applied this filter.

**Figure 2:**
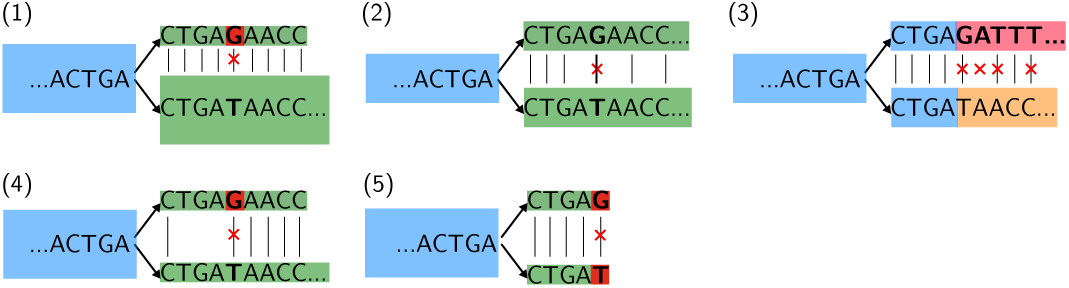
Unitig coverage is shown using node thickness. (1) A tip created by an erroneous ‘G’ in the De Bruijn graph. (2) Very similar patterns can be encountered with SNV (here G/T subsitution), however most of them are expected to create bubbles and to have a higher coverage. (3) Alternative variant differ from SNV by the larger Hamming distance. (4) Error in a shallow coverage region. (5) “Slingshot” error that creates more than one dead-end tips of low coverage.

In genomics, since most sequencing errors create bulges (pattern (3) in Figure 1) and tips (pattern (1) in Figure 2), their removal is done by a very efficient cleaning [1]. However, those patterns are commonly found as variants in transcriptomic De Bruijn graph [11], thus we introduce more advanced strategy to distinguish sequencing errors from actual variants in the next paragraph.

#### Advanced filters

In this section we present a likelihood model based on *k*-mer counting and hamming distance to attribute unitigs a probability to have emerged from an error or from a real variant. We use the Bayesian estimation framework to derive rules used to keep or discard unitigs for the correction step. Despite being based on simple framework, we demonstrate that these rules helps our correction to be more accurate than Bcool’s cleaning algorithm.

##### Preliminary observations We examine branching patterns

We start from the following observations: first, sequencing errors provoke patterns including a branching node and alternative paths in the graph. Thus we can focus on graph regions that correspond to triplets of nodes *u,v,w*∈*N* such that (*u,v*),(*u,w*)∈*E* or (*v,u*),(*w,u*)∈*E*. In the following, *u* is the branching node, *v* and *w* the alternative unitigs. In Figure 2, scenario (1) represents such branching pattern.

##### We examine pairs of alternative nodes and assume that the most covered comes from a real event

Second, for triplets *u,v,w*_*e*_, unitigs containing sequencing errors *w*_*e*_ are expected to be such that the coverages 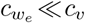 and 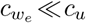. However, due to the skewed *k*-mer distribution, an error can be more covered than other low expressed real events in other parts of the graph. This motivates to work at a local scale (i.e. each branching pattern must be considered rather than applying a global threshold). Third, we expect that, for a triplet *u,v,w*_*e*_, if *w*_*e*_ is an erroneous unitig, *v* exists with 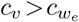 and corresponds a real event node. The existence of the error node without a true alternative is considered highly unlikely, unless in a specific scenario depicted in Figure 2 (5).

##### Sequencing errors have a small distance to real events

Within triplets where *w*_*e*_ is erroneous, we expect that *d*_*hamming*_(*v,w*_*e*_)*<t*, with *t*≤1%×|*w*_*e*_| because we consider a small (0.01% to 1%) and uniform error rate. On the contrary, alternative splicing or insertion/deletion for instance are expected to produce distant nodes (see Figure 2 (3) and (4) for the contrast). A SNV can yield a node with a small distance just as sequence errors. Such properties were also noticed and used in [3]. This means that in some cases, SNV with very shallow coverage might be lost during the correction.

##### Rules for the relative filters

We place ourselves in a given branching pattern with a triplet of nodes *u,v,w*, as described above (with *c*_*v*_ *c*≥_*w*_) and we want to determine whether *w* comes from an expressed variant or an error.

According to Bayesian frameworks [9] designed for close problems (to distinguish genomic variants), we consider the probability to observe an error *err* in a node given the data in the given pattern and the probability to observe a real event *event* under the same circumstances:

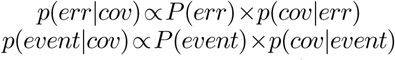

*cov* represents the observed data, i.e. the mean coverage in *k*-mers of each unitigs.

The first posterior probability corresponding to errors can be written as:

- *p*(*err*) = *ϵ*^*d*^, the probability to have errors in *w* (under the hypothesis that errors are uniformely distributed and that there is no error in *v*)
- 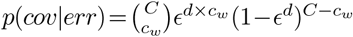, which is the probability to generate *c*_*w*_ erroneous unitigs given that the neighborhood indicates there are *C* unitigs (with *C* = *max*(*c*_*v*_ +*c*_*w*_,*c*_*u*_)).

The second posterior probability corresponding to an existing variant is:

- 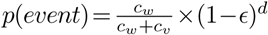 thus sequences are expressed and contain no sequencing error
- 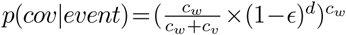 using the approximation of the expression of the region from which the unitigs originate, with their coverages as proxy

For each pair of unitigs, we compute both *p*(*err*|*cov*) and *p*(*event*|*cov*) and keep the maximum value between the two. If *p*(*err*|*cov*)>>*p*(*event*|*cov*) the unitig is removed, else it is considered as an event and kept in the graph for correction.

##### Implementation details

A detailed paragraph summing up the implementation details is provided in the Appendix. We mention that we apply an iterative procedure to compare pairs of nodes in case the branching node *u* has more out-neighbors. We also apply a low-pass filter on the *v,w* pairs to account for putative over-estimation of the error rate. Finally, polyA regions (from messenger RNAs) of the reads are removed before the graph construction to avoid highly connected nodes due to repeats.

#### Read correction

During the graph cleaning, some rare events may be removed from the graph due to their low coverage (example (2) in Figure 2). To cope with this peculiar situation that is not expected in genomic graphs, we developed a new read mapping algorithm. On such cases, our aim is to produce a local alignment of the read to the graph able to partly correct the read without applying incorrect modifications to the read subsequence absent from the graph.

#### Global strategy

The mapping procedure follows a seed and extend paradigm as described in [16, 17]. During the mapping of a read, the anchors shared between the read and the graph are detected and the alignment is greedily extended from each anchor using a simple linear scoring function, and the best alignments are used for correction. The proposed mapping algorithm brings two new important features described in detail below, the local alignment and the coverage assisted consensus correction.

#### Local alignment

Because we expect some read parts to be missing in the cleaned graph, we allow a trimming of the last (respectively first) unitigs selected if it improves the read score in order to find the best mapping possible, even if it does not involve the entire read. The procedure (algorithm described in Appendix) selects the unitig sequence optimizing the alignment score and keeps the corresponding unitigs for correction. Note that if a local alignment is chosen to correct a read, the unmapped part of the read will be unchanged. The rationale behind this choice is that we do not want to take the risk to perform wrong corrections in such cases (Figure 3).

**Figure 3:**
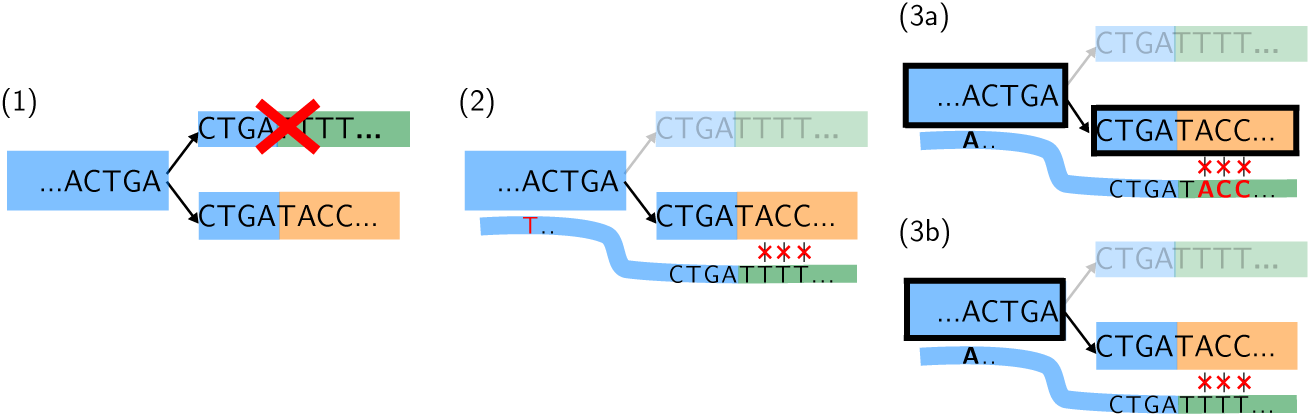
Partial correction strategy post mapping. (1) Before the mapping step, a real unitig is cleaned out from the graph because its coverage is too low. Henceforward in (2), reads from this region can only be mapped to the alternative path, leading to mismatches. In this example we present a read with a sequencing error (T in red) in its left part. Strategies consist in either not correcting the read at all, or fully correcting it (3a), leading to correct T but also to introduce many false positives on the right (unitigs used for the correction have a black stroke). (3b) Our strategy in BCT. If a a unitig bring a negative score, then this unitig is not considered for the correction leading to a partly mapped read. Then the best-scored among all possible mappings is chosen (other possible mappings are not shown in the example). Assuming the example mapping is the best-scored, this leads in the example to correct only the left part (T) of the read. This way, even if low covered parts of the graph were removed at the cleaning step, errors are not transmitted to next steps of the pipeline.

#### Coverage-assisted correction

Another significant feature of the mapping algorithm is its ability to take into account the unitigs coverage. Following the idea of Rcorrector that estimates the coverage of a read from its *k*-mers counts, our approach is able to estimate a read coverage using the coverages of unitigs it can be mapped on. This information is then used when different mappings present the same score, the average coverage of the read is computed from the unitigs shared among the mapping candidates. This behavior allows to choose more precisely the path that corresponds to the read context (an example is given in Figure 4, see also the algorithm in Appendix). Once again the rationale is to avoid to perform wrong corrections when possible. We use the statistics of the comparison of means test to decide whether one mapping should be selected among others or the corresponding read part should be left uncorrected (Implementation details in Appendix).

**Figure 4:**
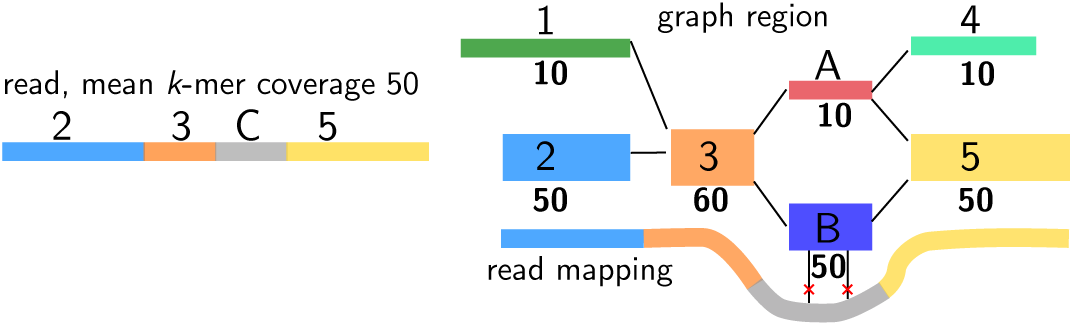
Choice of the most likely coverage in paths in case of a tie during mapping. Assuming a read containing a region C that can be mapped with a tie to two alternative unitigs A and B in the graph. The mapping strategy will take into account the most likely path to choose according to the possible coverages (bold numbers under the unitigs) in comparison to the coverage in *k*-mers of the read (here 50).

#### Differences with the other correction strategies

As BCT, Bcool and Rcorrector rely on De Bruijn graphs for the reads correction. Bcool was designed to correct genomic data with high uniform sequencing coverage. Its approach, based on a global alignment of the read to the graph is not adapted to RNA-seq data. Indeed, the reference graph may not contain all rare alternative events. Furthermore the graph cleaning step proposed by Bcool strongly relies on a uniform coverage.

Rcorrector uses paths of the De Bruijn graphs built from the reads’ *k*-mers for the correction. However, it does not perform a graph cleaning step. It selects *k*-mer nodes that are more covered than a threshold to correct reads in a greedy way. The correction step highly relies on the coverage estimation of the read to be corrected. Contrary to BCT, the graph is not compacted, thus each single *k*-mer coverage is used, and the filter threshold is estimated empirically.

Interestingly, proposed improvements over Bcool do not impact the ability of BCT to correct genomic datasets. We show in Appendix that BCT provides results similar to Bcool on the correction of a genomic dataset, with the same parameters.

## Results

### Transcriptomic De Bruijn graph cleaning

In this section we assess the behavior of our proposed method to remove erroneous *k*-mers from a De Bruijn graph constructed from non uniform dataset as RNA-seq data. To assess our ability to correct sequencing errors, we simulated RNA-seq dataset of more than 100 millions reads with FluxSimulator [12]. The reads were generated from RefSeq reference genome v19 and represent 107 millions reads, and 19 billions bases and were simulated with a one percent error rate. We selected rnaSPAdes [3] as it shares similarities with the presented approach (graph topology and hamming-based cleaning), the graph correction module of Bcool [17] to benchmark our own BCT implementation. We also included how a simple *k*-mer spectrum technique would perform for comparison.

In Table 1, we compared the produced cleaned De Bruijn graphs to the reference De Bruijn graph constructed using the errorless reads and measured the number of true positive (“good” *k*-mer seen both in the graph and the reference), false positive (“erroneous” *k*-mer in the graph not present in the reference) and the number of false negative (“missing” *k*-mer from the reference not found in the graph) and derive precision, recall and F1-score.

**Table 1:**
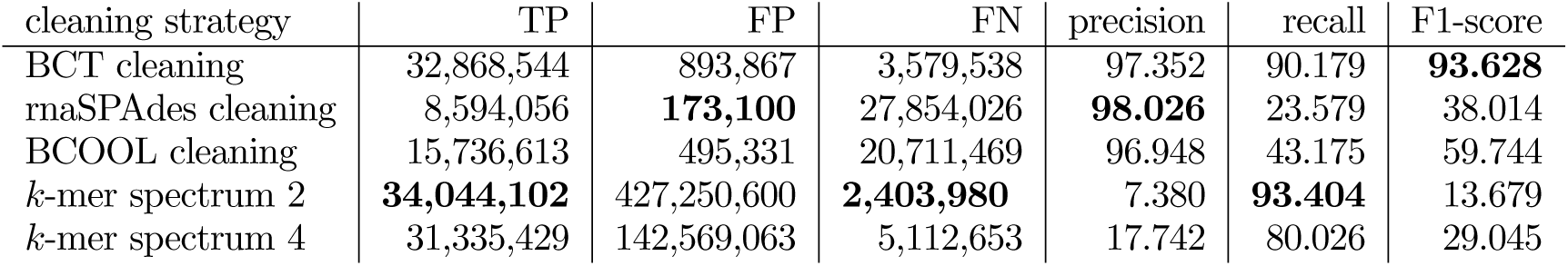
Evaluation of the De Bruijn graph cleaning step. True positive: TP, “good” *k*-mer seen both in the graph and the reference), false positive (FP, “erroneous” *k*-mer in the graph not present in the reference) and the number of false negative (FN, “missing” *k*-mer from the reference not found in the graph). We therefore define the recall and the precision of a correction as 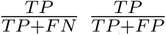 respectively. The F1-score is the harmonic mean of the recall and the precision. *k*-mer spectrum 2 and 4 strategies refer to the *k*-mer minimum abundance threshold: respectively 2 and 4.

We show that BCT’s cleaning is able to retain most *k*-mers, even *k*-mers with very low abundance. BCT lost 3.5 millions *k*-mers, slightly more than the filter removing unique *k*-mers but less than the filter removing *k*-mers seen 3 times or less. We observe that if rnaSPAdes or Bcool’s cleaning steps are able to suppress most sequencing errors (low FP and high precision) they tend to suppress a very important part of the expressed *k*-mers. In addition to its high recall BCT is also able to propose a similar ratio of erroneous *k*-mers than those methods.

### Results on simulated transcriptomic data

We kept the previously described dataset to assess our ability to correct sequencing errors. We corrected the read set using BFC [15] and Musket [18], that are correctors among the best performers according to recent benchmarks [17], Bcool and Rcorrector [27]. In Table 2, we report the recall and precision of our correction, the F1-score, the correction ratio (by which factor the error rate was divided after correction) and the proportion of reads that still contain at least one error after correction.

**Table 2:** Evaluation of the correction of simulated RNA-seq data. After correction we reported the number of true positive (TP),errors corrected to the right nucleotide, false positive (FP), nucleotides modified by the corrector to a wrong nucleotide and false negatives (FN), errors not corrected by the correctors. Precision, recall and F1-score are defined as in Table 1. The correction ration is defined as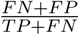. All corrector were used with default parameters as experimental evaluations showed a low impact of the parameters on the performances (See Appendix).

| corrector | precision | recall | F1-score | correction ratio | % erroneous reads |
| --- | --- | --- | --- | --- | --- |
| BCT | 99.532 | **98.511** | **99.019** | **51.248** | **2.069** |
| Rcorrector | **99.868** | 91.131 | 95.300 | 11.126 | 7.090 |
| Musket | 99.207 | 77.361 | 86.933 | 4.302 | 21.359 |
| BFC | 95.662 | 41.027 | 57.425 | 1.673 | 51.074 |
| Bcool | 84.856 | 25.002 | 38.624 | 1.299 | 64.5977 |

We observe that BCT is able to correct more errors than Rcorrector at the cost of being slightly less precise. We also observe that *k*-mer spectrum techniques developed for genomics data does not perform well.

Because of low expression variants, we expect the correction to be harder with a lower coverage as it become more challenging to distinguish erroneous from rare *k*-mer. To assess this effect we constructed smaller datasets by random sub-sampling on the one presented. We build a “medium” dataset of 53 millions reads and 9 billions bases, a “small” dataset of 17 millions reads and 5 millions bases and a “shallow” dataset of 13 millions reads and 2 billions bases. We kept Rcorrector as the most competing tool for this benchmark and report it in Table 3.

**Table 3:** Evaluation of the correction of simulated RNA-seq data according to the available coverage.

| dataset | corrector | precision | recall | F1-score | correction ratio | % erroneous reads |
| --- | --- | --- | --- | --- | --- | --- |
| Medium dataset<br>""" | BCT | 99.572 | **98.530** | **99.048** | **52.818** | **1.919** |
|  | Rcorrector | **99.851** | 91.153 | 95.304 | 11.134 | 7.083 |
| Small dataset<br>""" | BCT | 99.511 | **98.285** | **98.894** | **45.500** | **2.156** |
|  | Rcorrector | **99.827** | 91.007 | 95.213 | 10.929 | 7.223 |
| Shallow<br>""" | BCT | 99.442 | **97.692** | **98.559** | **35.015** | **2.697** |
|  | Rcorrector | **99.785** | 90.545 | 94.941 | 10.364 | 7.643 |

We show that BCT is able to increase its correction performances when a higher coverage is available. The correction ratio going from 35 from the 13 millions reads dataset to 52 with the dataset containing four time more reads. This can be explained that a higher coverage leads to the construction of a better De Bruijn graph as the sequencing errors and the variant become easier to distinguish. Per contra the performances of Rcorrector remained very similar despite an order of magnitude coverage change.

### Results on simulated metagenomic data

To evaluate the capacity of BCT to correct sequences from metagenomic samples, we created 2 synthetic datasets. First, we created a highly covered dataset (more than 100X) of 25 mixed bacterial genomes using CAMISIM [8] (genomes included in CAMISIM). Second, we created a low coverage dataset (20 in average) with high diversity of bacterial genomes (100 genomes). Datasets used for the reads generation can be found in Appendix. For both of the datasets, we added 1% of uniformly distributed errors.

Results are presented in Table 4. A study of the influence of BCT’s parameters is available in Appendix. While BCT, BFC and Rcorrector could handle 25 genomes correction, only BFC and BCT’s performances continued to keep up with the number of genomes. With default parameters, BCT had the best results on all metrics on the 100 genomes dataset.

**Table 4:** Evaluation of the correction of simulated metagenomic data. CAMISIM 25 genomes simulation contains 22,222,221,750 nucleotides split into 148,148,145 reads. 100 genomes simulation contains 7,654,414,260 nucleotides split into 51,023,830 reads. The average genome coverage is 20 with a lognormal distribution.

| corrector | precision | recall | F1-score | correction ratio | % erroneous reads |
| --- | --- | --- | --- | --- | --- |
| <u>CAMISIM 25 genomes</u> |  |  |  |  |  |
| BCT -S4 | 99.576 | **99.614** | **99.595** | **123.454** | **0.564** |
| Rcorrector | **99.953** | 99.233 | 99.592 | 123.001 | 0.628 |
| BFC | 93.560 | 66.218 | 77.550 | 2.726 | 37.786 |
| <u>100 genomes simulation</u> |  |  |  |  |  |
| BCT | **99.527** | **97.091** | **98.294** | **29.68** | **3.141** |
| Rcorrector | 97.745 | 48.986 | 65.265 | 1.919 | 41.07 |
| BFC | 98.943 | 95.686 | 97.287 | 18.759 | 3.951 |

### Results on real datasets

We tested our correction on more realistic scenarios from human sequencing: 2 RNA-seq files and 5 single cell RNA-seq (scRNA) files, presenting various read sizes and coverages. Since no ground truth is available when working on real data, we used read alignment on the reference genome as a proxy to correctors’ performances. All read files (raw and corrected) were mapped to the human reference genome using STAR [7] with default parameters. Since BayesHammer uses different information (FASTQ qualities) than our algorithm, we do not report direct comparison. However, BayesHammer’s results can be found in Appendix, showing that it performed best on scRNA-seq, with BCT ranking second in terms of mismatches. Results are presented in Tables 5 and 6, and the complete results are available in Appendix. BCT successfully reduced the mismatch rates, at a better rate than Rcorrector in 6 out of 7 cases. It increased the reads length mapped and decreased the fraction of unmapped reads in most cases.

**Table 5:** Correction of RNA-seq human datasets. Statistics were computed using STAR 2.7.0.

| metric | SRR3466506 |  |  | SRR8942950 |  |  |
| --- | --- | --- | --- | --- | --- | --- |
|  | raw | BCT | Rcorrector | raw | BCT | Rcorrector |
| average mapped length | 199.29 | 199.43 | 199.30 | 99.93 | 100.03 | 99.97 |
| unmapped (%) | 0.69 | 0.80 | 2.3 | 3.08 | 3.05 | 3.46 |
| mismatch rate per base (%) | 0.31 | 0.18 | 0.30 | 0.44 | 0.33 | 0.39 |

**Table 6:** Correction of scRNA-seq human datasets. Statistics were computed using STAR 2.7.0.

| metric | ERR2457632 |  |  | ERR2457650 |  |  | SRR2977735 |  |  |
| --- | --- | --- | --- | --- | --- | --- | --- | --- | --- |
|  | raw | BCT | Rcorrector | raw | BCT | Rcorrector | raw | BCT | Rcorrector |
| average mapped length | 71.92 | 73.51 | 73.53 | 71.35 | 73.44 | 73.37 | 43.59 | 43.74 | 43.78 |
| unmapped (%) | 7.9 | 7.81 | 7.92 | 11.76 | 11.74 | 12.14 | 0.45 | 0.57 | 1.06 |
| mismatch rate per base (%) | 0.45 | 0.27 | 0.27 | 0.76 | 0.49 | 0.60 | 0.71 | 0.62 | 0.68 |
|  | SRR5739564 |  |  | SRR4055098 |  |  |  |  |  |
|  | raw | BCT | Rcorrector | raw | BCT | Rcorrector |  |  |  |
| average mapped length | 96.26 | 96.27 | 96.47 | 71.41 | 71.44 | 71.62 |  |  |  |
| unmapped (%) | 37.29 | 37.16 | 37.39 | 0.61 | 0.7 | 0.65 |  |  |  |
| mismatch rate per base (%) | 0.38 | 0.24 | 0.25 | 1.19 | 1.21 | 1.03 |  |  |  |

### Conclusion and future works

In this work we presented a method able to construct a De Bruijn graph containing a low amount of erroneous *k*-mers while preserving most of the diversity present in dataset, even if in presence of a high discrepancy in the observed coverage. We shown that this graph could be used as a reference for read correction using De Bruijn graph local read alignment. We benchmarked the implementation of the proposed method, dubbed BCT, against state-of-the-art correctors on RNA-seq and metagenomic data and showed the interest of such a method to efficiently raise the signal to noise ratio in such datasets.

Since our proposition is designed as a workflow, a natural extension of this work could be to test different graph alignment tools as graphAligner [19] or vg [10] to assess if they could be adapted to perform read correction.

An ongoing application of this work is the hybrid correction of long reads. Transcriptomic long reads (Iso-seq and Nanopore) are increasingly used for full-length isoform identification and *de novo* transcript and alternative splicing discovery. For these data, hindered by high error rates and especially insertion/deletion, a primary correction step is often required. The mixture of short and long reads was shown to provide sequences with higher accuracy [6], in particular for technologies prone to systematic errors such as Oxford Nanopore. We would like to investigate the impact of high quality short reads on such correction.

Another potential function of the proposed method would be to produced paired-end read alignment to produce merged long reads from read pairs [4]. Such extended and corrected reads could improve assembly or variant calling, especially in repeated regions. Those extended reads could also improve the hybrid correction as they should be easier to associate to a long reads.

An interesting feature of our method is that most reads can be decomposed into a path of the De Bruijn graph. This compressed representation could be highly efficient to index RNA-seq experiment at the read level. One could build an index of such paths to efficiently associate a *k*-mer (or a set of *k*-mers) to the reads it originates from.

Finally, we see in Figure 5 that the corrected reads account for orders of magnitude less distinct *k*-mers to index while preserving most of the original read *k*-mer content. Therefore, this procedure could greatly impact the scalability of *k*-mers based indexing methods such as SBT [26] or BIGSI [2] by drastically reducing the amount of *k*-mers to index.

**Figure 5:**
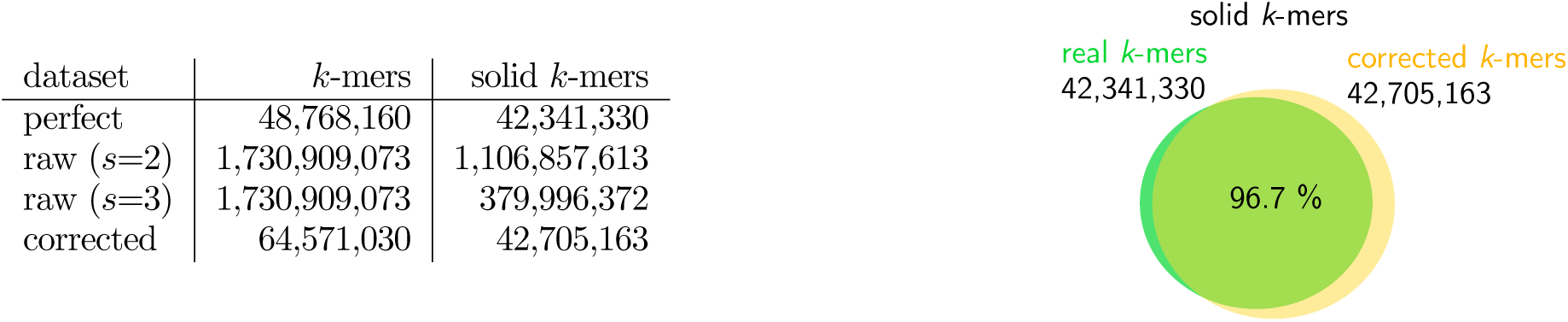
Effect of the correction on the usable *k*-mer set size.The simulated dataset from Table 2 was used. The perfect dataset refers to reads without errors. The corrected dataset was generated with BCT. We filtered *k*-mers with abundance ¡ 2 to obtain solid *k*-mers for these two datasets. We show *k*-mers from raw reads, with abundance filters ¡ 2 and ¡3 for solid *k*-mers. On the right, the proportion of solid corrected *k*-mers that intersect real *k*-mers is presented.

## Acknowledgments

The authors would like to thank Rayan Chikhi and Mikaël Salson for their support and the insightful discussions. This work was supported by the ANR Transipedia (ANR-18-CE45-0020). We also thanks the Universitè de Lille HPC Cloud computing resources and the Institut Pasteur for the computing and storage resources.

## Footnotes

1 number of reads that cover a given nucleotide in the original sequences

